# Efficacy of functional connectome fingerprinting using tangent-space brain networks

**DOI:** 10.1101/2025.01.17.633606

**Authors:** Davor Curic, Sudhanva Kalasapura Venugopal Krishna, Jörn Davidsen

## Abstract

Functional connectomes (FCs) are estimations of brain region interaction derived from brain activity, often obtained from functional Magnetic Resonance Imaging recordings. Quantifying the distance between FCs is important for understanding the relation between behaviour, disorders, disease, and changes in connectivity. Recently, tangent space projections, which account for the curvature of the mathematical space of FCs, have been proposed for calculating FC distances. We compare the efficacy of this approach relative to the traditional method in the context of subject identification using the Midnight Scan Club dataset, in order to study resting-state and task-based subject discriminability. The tangent space method is found to universally out-perform the traditional method. We also focus on the subject identification efficacy of subnetworks. Certain subnetworks are found to outperform others, a dichotomy which largely follows the ‘control’ and ‘processing’ categorization of resting state networks, and relates subnetwork flexibility with subject discriminability. Identification efficacy is also modulated by tasks, though certain subnetworks appear task independent. The uniquely long recordings of the dataset also allow for explorations of resource requirements for effective subject identification. The tangent space method is found to universally require less data, making it well suited when only short recordings are available.

**AUTHOR SUMMARY:** Functional connectomes, which describe the similarity between the recorded activity of different brain regions, are ubiquitous for researcher trying to understand brain dynamics on short and long time scales in the presence and absence of neurophysiological diseases. This work applies a Tangent Space approach, a novel method to calculate the distance between functional connectomes, to a unique high-quality dataset (the Midnight Scan Club) in order to better understand the variability and uniqueness of connectomes across subjects, and how subject identification (also called fingerprinting) compares across tasks. We also show that not only does the tangent space method offer greater sensitivity to changes, it does so with significantly fewer resources, and in some scenarios reveals qualitatively different interpretations than the traditional method.

## INTRODUCTION

Functional connectivity is a mathematical estimate of the interactions between different brain regions, often derived from pairwise correlations (or other similarity measures) of neurophysiological signals such as BOLD signals obtained in fMRI scans (Bastos & Schoffelen, 2016; Martin et al., 2017; Oliver, Hlinka, Kopal, & Davidsen, 2019). In contrast to structural brain connectivity derived from physical connections of neurons or white matter tracts (Griffa, Baumann, Thiran, & Hagmann, 2013; S^̌^koch et al., 2022), functional connectivity depends on the context (e.g., rest, task, etc.) and can change on very short time scales. Combined with tools from network neuroscience (Bassett & Sporns, 2017), **functional connectomes** (FCs), the matrix representation of functional connectivity, have been used to study group-averaged properties of the brain in the resting state, during tasks, and in the context of disease (Allen et al., 2022; Cabral, Kringelbach, & Deco, 2014; Du, Fu, & Calhoun, 2018; Faryadras, Burles, Iaria, & Davidsen, 2024). Increasingly there is also interest in applying FCs in clinical settings, and as such, an accompanying interest in studying single-subject variability for both structural and functional connectivity (Allen et al., 2022; Barch et al., 2013; Gordon, Laumann, Gilmore, et al., 2017; Mueller et al., 2013; Oliver et al., 2019; S^̌^koch et al., 2022). This is on top of the already-existing interest in how different anatomical regions of the brain can functionally vary across individuals (Kane & Engle, 2002; Vogel & Machizawa, 2004).

To this end, it has also been demonstrated that the subject-specific differences of FCs can serve as a ‘fingerprint’ for identifying individuals (Finn et al., 2015), thus motivating FC **fingerprinting**. The original formulation is as follows; given a known database of FCs belonging to unique individuals, and a unknown ‘target’ FC, the task of fingerprinting is to identify which individual in the known database the target FC belongs to, with the identification accuracy being calculated as a measure of how many correct identifications are made. Using this approach, Finn et al. (2015) were able to demonstrate identification accuracies of 94% when using whole brain FCs. More recently, the notion of fingerprinting has been expanded beyond the task of subject identification, moving towards applying it in clinical settings. For example, patients with schizophrenia exhibited less longitudinally stable FCs relative to healthy controls (Kaufmann et al., 2018), while other studies have applied fingerprinting to identify predictors of treatment responds Nemati et al. (2020). We also point the reader to a recent review of fingerprinting analysis (Ramduny & Kelly, 2024).

Ultimately the ability and efficacy of distinguishing individuals, or the same individual longitudinally, comes down to a question of “distances” — FCs belonging to the same subject should have lower distances in some mathematical sense which can be used as a classifier (Fawcett, 2006). Temporally diverging FCs could then also indicate the presence and evolution of disease (Supekar, Menon, Rubin, Musen, & Greicius, 2008), or perhaps converging FCs could reflect the efficacy of treatment for disorders such as depression (Wang, Hermens, Hickie, & Lagopoulos, 2012). Accurately defining distances in FC space can also be important to understand dynamical functional connectivity on short timescales and how it relates to behaviour (Menon & Krishnamurthy, 2019).

However, defining a distance on the mathematical space of FCs is not trivial. FCs generated from correlation analysis are **positive semi-definite** (meaning all eigenvalues are greater than, or equal to, zero) (Wasserman, 2013). This imposes constraints on the possible values that the elements of the matrix can take on, which in turn curves the space of possible FCs, creating a manifold of possible values called the **positive semi-definite manifold** (Bhatia, 2015). For covariance matrices this space is an infinite convex cone (Abbas et al., 2021). For correlation matrices (i.e., the time series is first standardized by z-scoring) the space is a **compact convex manifold** (Rousseeuw & Molenberghs, 1994). Traditional measures of distance (or equivalently similarity) like the correlation distance, do not take into account the curvature of the space of FCs. Analogously to measuring the separation between cities on the surface of the Earth with a straight line in three-dimensions, metrics that do not take into account the curvature of the space of positive semi-definite matrices can mischaracterize distances.

**Geometry-aware methods** are a set of techniques that take into account the curvature of this space (Venkatesh, Jaja, & Pessoa, 2020). One such method is the **tangent space** (TS) method which involves projecting FCs onto a flat space that is tangent to a point on the surface of the convex manifold (Abbas et al., 2023; Ng, Varoquaux, Poline, Greicius, & Thirion, 2016). Once in this tangent space, traditional metrics of distance can be freely applied as this new space has zero curvature. Such an approach has been shown to improve the sensitivity of subject identification over traditional methods. For example, the work presented in (Abbas et al., 2023) shows a nominal identification rate of monozygotic twins of 0.5 when using the non-TS method, and perfect identification when using the TS method (note however this analysis used a modified version of the test-retest method presented in (Finn et al., 2015)). Importantly, they also observed that when distances in Tangent Space are calculated using the correlation similarity, the identification rates are not only high (often near or at unity and always higher than the non-TS method), but are also effectively independent of several hyperparameters (parcellation and regularization), reducing the amount of hyperparameter tuning relative to other geometry-aware approaches like the geodesic distance (Abbas et al., 2021; Venkatesh et al., 2020).

Here we build on prior work and apply the tangent space method to the Midnight Scan Club (MSC) dataset (Gordon, Laumann, Gilmore, et al., 2017) (10 subjects, 5 male, 5 female between 24 and 34 years old (average 29.1 ± 3.3 years)). The MSC dataset was chosen as its unique features complement prior studies (Abbas et al., 2023; Finn et al., 2015). Specifically, the MSC dataset is part of a growing effort by researchers to precisely study the per-subject variability of the functional connectome, motivated in part by improving individual-level applications of FCs to clinical settings (Gratton et al., 2020). To this end, the MSC dataset contains five hours of resting state data, and six hours of task-based data for each subject, compiled over ten sessions. The uniquely long sessions offered by the MSC dataset allow for deep explorations of inter-subject variability which can expand prior studies in FC fingerprinting (Abbas et al., 2023; Finn et al., 2015). In particular, we explore how different subnetworks contribute towards subject discriminability, including to what extent subject discriminability is task dependent. If there is variation in identification efficacy between subnetworks, what is the connection between strongly performing and poorly performing subnetworks (Vanderwal et al., 2017)? For example, previous studies have suggested a dichotomy of ‘control’ and ‘processing’ subnetworks which play different roles in the context of tasks, and have different network theoretic properties (Power et al., 2011). Complementing this are observations of greater flexibility in certain subnetworks like the frontopariatal subnetwork relative to unimodal sensory areas (e.g., visual, auditory) (Mueller et al., 2013; Vanderwal et al., 2017). While prior work has used non-geometry aware techniques to study the fingerprinting efficacy of subnetworks (Finn et al., 2015), here we take advantage of the increased sensitivity of TS analysis to gain further insight into the relationship between fingerprinting efficacy and inter-subject variability.

In this manuscript, we address the aforementioned questions and while we find in particular the dichotomy of ‘control’ and ‘processing’ subnetworks to be true for previously used (non-TS) methods, the tangent space approach suggests some deviations from it. We also compare the efficacy of the TS method against the traditional (non-TS) method (here the correlation distance) of subject discrimination to be able to contrast our findings against previous work (Finn et al., 2015), finding that the TS method universally outperforms the non-TS method. Additionally, while the large number of long scans per subject in the MSC dataset (ten for each category of recordings, spanning ten consecutive days) are important for answering fundamental questions regarding subject variability (Gordon, Laumann, Gilmore, et al., 2017), a natural question follows — is that much data actually required? Thus, we explore the resource requirements for subject identification (e.g., number of recordings, duration of recordings), again comparing the TS and non-TS methods, and find that the TS method requires substantially fewer resources to perform as well as the non-TS method. Moreover, the TS method offers extra degrees of freedom for performing the projection (i.e., selecting the point at which the projection is made). While previous work has studied the different mathematical techniques one can use to obtain the reference point (Abbas et al., 2023), it is not clear whether certain FCs (e.g., resting or task-based) perform better as a reference. We observe that subject identification is comparable across all tasks with regards to this, which lessens the number of pre-existing recordings a fingerprinting task would be required to have. Answering all these questions of practicality and implementation is particularly important for possible future applications.

## METHODS

### Dataset

Here we analyze the open source Midnight Scan Club Dataset, as described in greater detail in (Gordon, Laumann, Gilmore, et al., 2017). Briefly, the dataset consists of ten rest fMRI recordings (over ten subsequent days) per ten subjects; 5 male, 5 female between 24 and 34 years old (average 29.1 ± 3.3 years). Each session started with thirty minutes of resting state recording, which involved visually fixating on a white crosshair presented against a black background. Subsequently, each subject was then scanned during the performance of three separate tasks: motor (2 runs per session, 7.8 min combined), mixed design (2 runs per session, 14.2 min combined), and incidental memory (3 runs per session, 13.1 min combined). The subjects had to exhibit directed movement in response to visual cues in the motor task, discriminated between patterns and words in mixed task, and made binary decisions about faces, scenes and words in the memory task.

A Siemens Trio 3T MRI scanner was used to record fMRI data. Axial functional images were taken with the following specifications: TR = 2.2 s, TE = 27 ms, flip angle = 90°, voxel size = 4×4×4 mm^3^, matrix size = 64×64×36. A high resolution T1-weighted structural MRI recording (TR/TE/TI = 2300/3.7/1000 ms, voxel size = 0.8 × 0.8 × 0.8 mm^3^) was used as anatomical reference. One subject (MSC08) reported drowsiness, and thus was excluded from this analysis (Gordon, Laumann, Gilmore, et al., 2017). This left 90 resting state recordings (ten per subject, with nine subjects). The Gordon-333 atlas which parcellates the fMRI data into time series for 333 common regions of interest (ROI) (Gordon et al., 2014; *Midnight Scan Club: ROI-based information*, 2017) was used to ensure the size of each FC was the same across subjects (333 × 333), a requirement of this analysis. Previous work has shown that identification rates when using the correlation distance, which we use in this manuscript (discussed below), is optimal and independent of parcellation granularity beyond 100 parcels (Abbas et al., 2023), and thus this ‘hyperparameter’ is fixed to 333 in this manuscript.

The preprocessing pipeline involved reducing artifacts, maximising cross-section registration and distortion correction by mean-field mapping. For resting state recordings, periods of motion were removed with the provided temporal masks. These were identified as volumes above a frame-by-frame displacement (FD) threshold of 0.20 mm, as per (Power et al., 2014). Frames with FD *>* 0.2 mm were labeled as motion-contaminated and removed from the recording. The total amount of removed frames was 28% ± 18% of the total recording time (Gordon, Laumann, Gilmore, et al., 2017). For task recordings, an additional fixation temporal mask was used to remove periods of no fixation (Gordon et al., 2023). Several task-based recordings did not have accompanying temporal masks, or had masks whose length did not match the length of the recording. These recordings were thus discarded. One subject (MSC09) had five sessions discarded and so this subject was not considered for the task analysis. As we wanted to maintain the same sessions across all subjects, any session that was not common to all remaining subjects was discarded. This resulted in 48 total recordings (six sessions per subject, eight subjects).

### Estimating functional connectivity matrices

Resting state and task-based FCs were constructed by calculating the pairwise zero-lag correlation coefficient. For each pair of regions *m, n*, the correlation coefficient of their associated time-courses *v_m_*(*t*) and *v_n_*(*t*) is calculated as

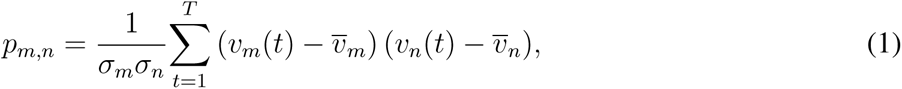

where *T* is the total number of samples after removal of motion and fixation artifacts, *σ_x_* is the standard deviation of *v_x_*, and *v_x_* is the average of *v_x_* over the *T* time points. This results in a 333×333 symmetric positive semi-definite matrix **P** for each condition in a recording session of a given subject. An example of an FC is given in Fig. 1a.

**Figure 1.**
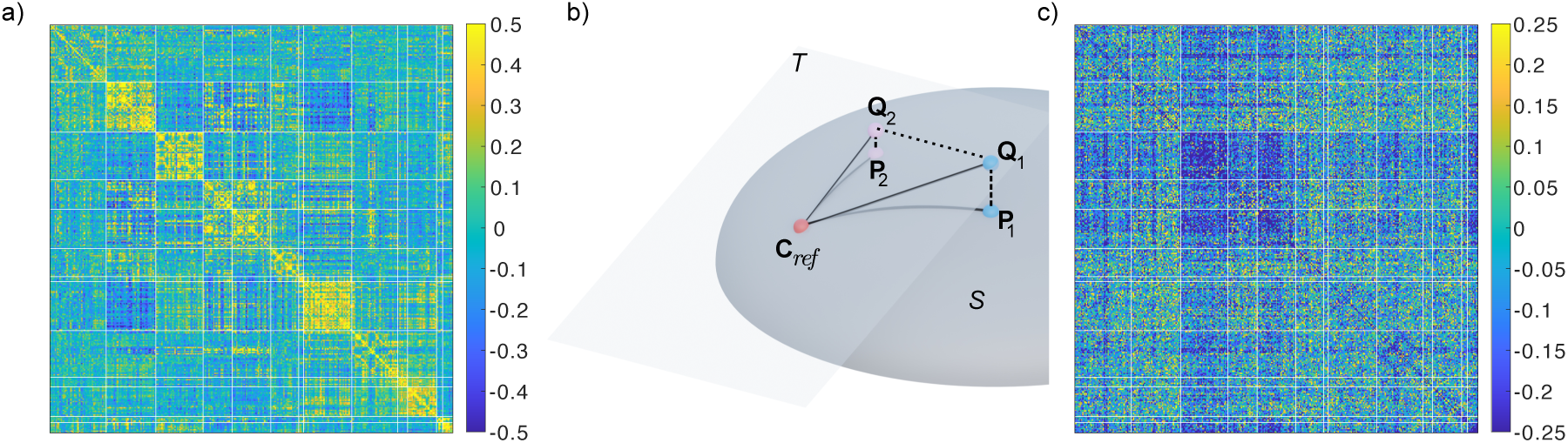
a) Functional connectivity (FC) matrix for a single resting state recording. Rows and columns have been grouped according to which resting-state network the regions belongs to in the Gordon 333 atlas. Each resting-state network is highlighted by the thicker lines. b) Pictorial demonstration of tangent space projection. The FC matrices **P**_1_, **P**_2_ belong to a curved manifold of positive definite matrices, *S,* and thus distances between any of the matrices are non-Euclidean. Choosing **C**_ref_ as a reference point, the tangent space, *T*, is a new linear vector space tangent to *S* at the point **C**_ref_. The FC matrices **P**_1,2_ in *S* are projected onto the tangent space *T* where they are labeled as **Q**_1,2_, for which the distances are Euclidean. c) Tangent space representation of the FC shown in a) using the logarithmic mean over all resting state recordings as **C**_ref_ (see Eq. (4)).

### Tangent space projection of FC matrices

The set of positive semi-definite matrices comprises a convex manifold. For covariance matricies this manifold forms an infinite cone (Abbas et al., 2021), but for the correlation matrices (i.e., each signal is mean subtracted and divided by the respective standard deviation) the manifold is compact (Rousseeuw & Molenberghs, 1994). Euclidean measures of distance may assign significantly biased distances between pairs of FCs in the curved space. One approach to remedy this is to calculate the geodesic distance between pairs of FCs (Bhatia, 2015; Venkatesh et al., 2020):

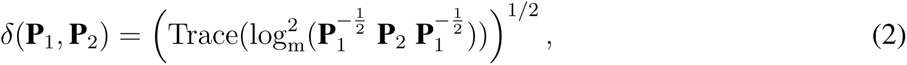

where log_m_ is the matrix logarithm (*logm*, MATLAB). However, the geodesic distance can be quite sensitive to the specific regularization used (Abbas et al., 2021). Another approach (the one used here) is to project FCs onto a space tangent to the manifold at around a **reference matrix C***_ref_*. Then one can use Euclidean distance metrics or related distance measures within this linearized space (Abbas et al., 2023; Ng et al., 2016).

Mathematically, for a given reference matrix **C***_ref_*, the projection of an FC **P** from the original positive semi-definite manifold to the tangent space is given by (Abbas et al., 2023; Bhatia, 2015; Ng et al., 2016),

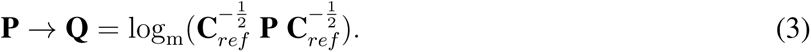

Note that for clarity we reserve the letters **P** to refer to matrices in the positive semi-definite manifold, and **Q** for their projected TS equivalents, with the exception of the reference **C***_ref_*. A demonstration of this procedure is depicted in Fig. 1b, with the resulting tangent space projected FC shown in Fig. 1c.

The choice of the reference matrix can impact the quality of the estimation of the distance between matrices. One choice would be to use the arithmetic average over all given FCs. However, previous studies have shown that alternative centroids perform better, such as the log-Euclidian and the Riemannian mean (Abbas et al., 2021; Fiori, 2009). Our analysis suggested the same with the log-Euclidean and Riemannian mean performing on par with each other. The log-Euclidean centroid is mathematically more straightforward and is thus the one presented here. For a set of FCs **P***^i^*, the reference can be calculated as

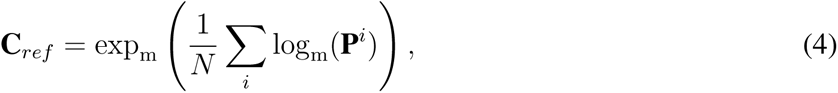

where exp_m_ denotes the matrix exponential (*expm*, MATLAB).

An important consideration for the tangent space projection (i.e., Eq. (3)) is the invertability of **C***_ref_*. This matrix is non-inevitable if det(**C***_ref_*) = 0. Because the determinant is equal to the product of all the eigenvalues, **C***_ref_* is non-invertable if there exists at least one zero eigenvalue. If this is indeed the case, **regularization** can be performed by adding *⋋***1** to **C***_ref_*, where *⋋ >* 0 and **1** is the identity of commensurate size as **C***_ref_* (Venkatesh et al., 2020). The effect of regularization raises the eigenvalues of **C***_ref_* such that **C***_ref_* + *⋋***1** is invertable. For clarity we focus predominately on results for *⋋* = 1, however also tested *⋋* = 0.1 and *⋋* = 10.

### Correlation distance between matrices

Given an arbitrary pair of matricies **X***^i^* and **X***^j^* (where **X** is a stand in for **P** or **Q**), fingerprinting is the process of determining whether or not these two correspond to the same individual based on some distance measure *d_i,j_*. Here, we use the correlation distance between the **X***^i^* and **X***^j^*. While strictly speaking *d_i,j_*is not a distance metric (e.g., it does not obey the triangle inequality), it has been shown to have several useful properties, namely being effectively independent of both choices in regularization as well optimal and parcellation independent beyond 100 parcels (Abbas et al., 2023), thus reducing several hyperparameter choices. Because **X** is symmetric the correlation distance *d_ij_*is calculated using only the *n*(*n* − 1)*/*2 upper-triangular elements of **X** (i.e., excluding the diagonal) (Abbas et al., 2023):

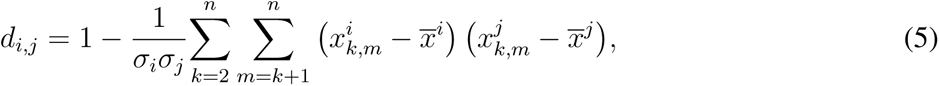

where 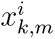 is the *k, m*-*th* element of **X***^i^*, and *x_i_* and *σ_i_* are the average and standard deviation across all *n*(*n* − 1)*/*2 elements of the upper triangle of **X***^i^*. We will denote **D** as the distance matrix indexed by all pairs *i, j*.

### Fingerprinting/Classification

The distance matrix **D** can be decomposed into block-diagonal form, where each block corresponds to a single subject, and the size of the block is the number of FC’s (or recordings) belonging to said subject. The elements of **D** that are within such a block are called intra-subject distances, while elements of **D** outside one of these blocks are inter-subject distances. For the MSC the number of inter-subject distances is 7200, while the number of intra-subject distances is 810. Because approximately 90% of the data belongs to the inter-subject class, the dataset is said to be imbalanced. We generate two cumulative distance distributions from the respective ‘within’ and ‘between’ subject values in **D**.

For any pair of FCs **X***^i^* and **X***^j^*, subject discrimination can be reduced to a binary classification problem (Géron, 2022); for some discriminating threshold 0 ≤ *θ* ≤ 2 (2 was found to be a a suitable upper bound for our dataset), **X***^i^* and **X***^j^* belong to the same subject only if *d_ij_ < θ*. For a given *θ* it is expected that there will be a number of true positive (TP) classifications, false positive (FP) classifications, true negative (TN) classifications, and false negative (FN) classifications. The accuracy (ACC) of the classification model can be calculated as the following:

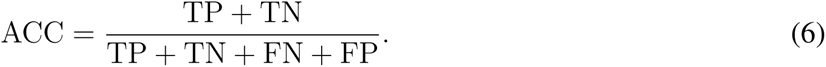

While the accuracy can be a useful metric, it can be misleading in imbalanced datasets. For example, a trivial classifier which assigns every FC as not belonging to any subject would perform with 0.9 accuracy. One approach to address this problem is to use the **Receiver Operating Characteristic** (ROC) curve (*prefcurve*, MATLAB) which plots the true positive ratio, defined by (Fawcett, 2006)

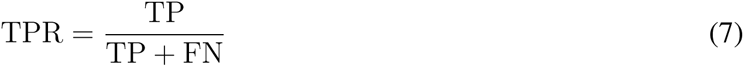

against the false positive ratio, defined by

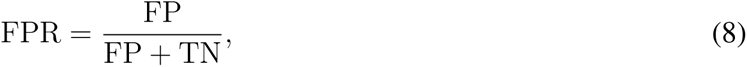

which are calculated across all values of *θ.* The ROC can be directly related to the cumulative distribution functions (CDF) of the inter- and intra-subject distances in the following way: for all values of *θ*, plot the value of the inter-subject CDF at *θ* against the value of the intra-subject CDF at *θ*. Perfect subject discriminability is possible if there exists a *θ* such that the intra-subject CDF evaluates at one, while the inter-subject CDF is simultaneously zero. This corresponds to a step-function in the ROC plot which starts at the point 0,0, and is otherwise unity, while total indiscriminately (i.e., chance level of subject discrimination) would be a diagonal line where TPR and FPR are equal. The **Area Under the Curve** (AUC) of the ROC is a commonly used metric to summarize the ROC. For a perfect classifier AUC= 1 where as AUC= 0.5 for chance levels of subject discrimination. In general, the closer the AUC is to unity the better the classifier.

An alternative metric for quantifying the quality of a classifier is to use the **Precision-Recall** (PR) curve which instead plots precision, TP/(TP + FP), against recall, TP/(TP + FN), across all thresholds *θ*. The PR curve has been found to be more robust in the context of imbalanced datasets (Saito & Rehmsmeier, 2015). The precision can be interpreted as the proportion of correctly classified pairs of matrices among all the positive predictions made by the classification procedure, while recall is proportion of correctly classified pairs among all the actual positive instances (Géron, 2022). The AUC is again a commonly used metric for summarizing the PR curve. However, unlike the AUC-ROC which is bounded between 0.5 and 1, the lower-bound of the AUC-PR is equal to the accuracy of a random chance classifier.

### Sampling shorter recording segments

To test the efficacy of fingerprinting with respect to the length of the recording we apply a boot-strapping approach. First, a random frame of the recording is chosen, and segments of length *l* are chosen on each side of the random point. The resulting window is of length 2*l* + 1, but we emphasize here that, because frames with large motion periods are removed, these frames may not have been originally sequentially adjacent. We then calculated the FC matrix from points within this window (Eq. 1), and repeated the above fingerprinting procedure. This random sampling was repeated. For large windows *l*, the probability to re-sample the same section of the recording is high, while for small windows re-sampling becomes less probable, thus the number of trials should depend inversely on *l*. We chose this to be ⌊*L/*2*l*⌋, where *L* = min(*T*_1_*, T*_2_*, …, T_N_*) is the smallest duration of all of the *N* recordings being considered (which is needed as artifact removal results in unequal total recording duration). For each trial the AUC is calculated and finally averaged across each trial, with the standard deviation for errorbars.

### Resting-state subnetwork analysis

The Gordon 333 atlas is subdivided into a set of 13 non-overlapping resting-state networks (RSN) (Adhikari et al., 2021). The size of these RSNs ranges from four nodes (salience network) to 41 nodes (the default mode network), with 47 nodes belonging to an ‘unassigned’ category. We found that fingerprinting using RSNs was unreliable for particularly small RSNs. As such we discarded all RSNs with less than 10 nodes (4 RSNs), leaving a total of 8 RSNs for the analysis: visual (VIS), somato-motor hand (SMH), auditory (AUD), cingulo-opercular (COP), ventral attention network (VAN), dorsal attention network (DAN), frontal parietal network (FPN), and default mode network (DMN). Once RSNs have been established, we generated FC matricies for each RSN separately. We then recalculated the distance matrix **D** and repeated the fingerprinting analysis using the TS and non-TS methods, and separability is calculated from the ROC for each case. This was done for both resting-state recordings as well as task-based recordings.

### Subject identification using a known database

In order to mimic how fingerprinting might realistically be employed in a clinical setting, we divide the collection of FCs into two groups; a database of *k* FCs per *N* subject for which the corresponding subject is known, and a target FC whose corresponding subject is not known (Finn et al., 2015). The pairwise distance between the target and each of the *Nk* database FCs is computed using both the TS and non-TS method. In the case of the TS method, this is done by calculating a reference point using all *Nk* database FCs. The *k* database FCs are chosen randomly from the 10 sessions, without replacement. A single target is then chosen from the remaining *N* × (10 − *k*) FCs.

To identify the subject of the target FC we use a majority rules system; we calculate the mean target distance, averaged over all *k* known subject FCs. We classify the target FC as belonging to the subject with the lowest average distance. A point is awarded only if the correct subject is identified. This is repeated 100 times, with a total score calculated as the percentage of the number of successes, which we compute for each *k*. We performed this analysis for the resting state data as well as the somato-motor subnetwork (see below) during the motor task as a worst-case scenario.

## RESULTS

From the MSC dataset, FCs are generated from pair-wise correlations of the parcellated regions, as described in Methods. One such FC is shown in Fig. 1a. We attempt to identify individuals based only off distances between these FCs using two methods; first, by projecting FCs into a tangent space where the distance is subsequently calculated (the TS method, see also Fig. 1b,c), and second, by calculating distances directly between the functional connectomes (non-TS), as described in Methods. In both cases the pairwise correlation distance (Eq. 5) serves as the distance metric. The identification rates of each method are then compared. We will first analyze the resting-state recordings, and then task-based recordings. Finally we analyze sub-networks of the Gordon 333 parcellation and the robustness of the results against recording length.

### Resting State

We focus first on the 90 resting state recordings (ten for each of the nine subjects). The reference point for the tangent space projection was chosen to be the logarithmic mean calculated across all recordings (Eq. 4), using *⋋* = 1 for regularization. The pairwise distances (for either TS or non-TS methods) were gathered into a distance matrix **D** (see Methods), shown in Fig. 2a. When sorted by subject, **D** is block-diagonal, with low intra-subject distances going along the main diagonal, and high inter-subject distances off the the diagonal. This indicates that functional connectomes belonging to the same individual (intra) are closer than those belonging to different individuals (inter), as expected (Finn et al., 2015; Gordon, Laumann, Gilmore, et al., 2017).

**Figure 2.**
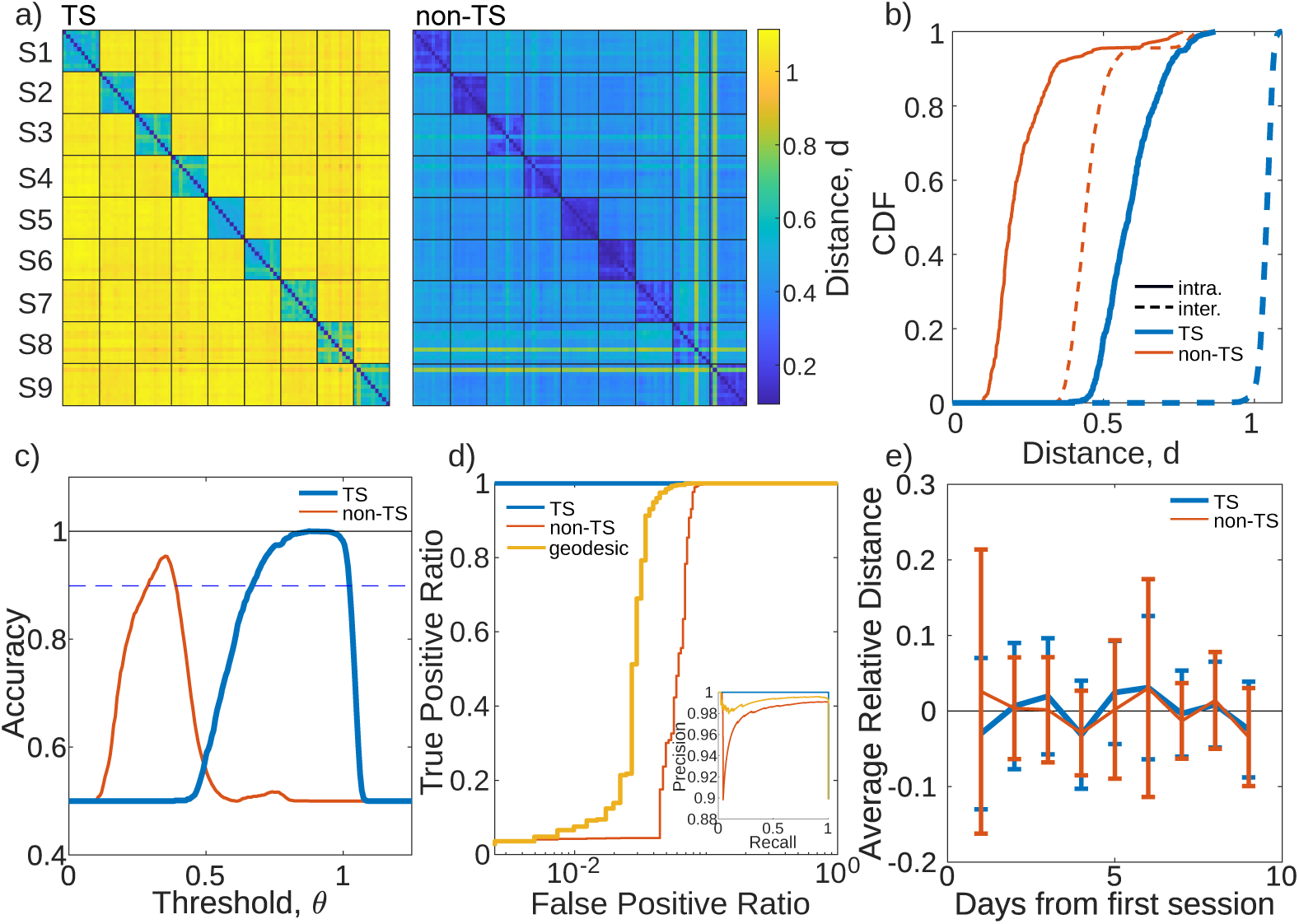
a) Pairwise correlation distance matrix between tangent space (TS) projected FCs (left) and non-TS FCs (right) for the nine subjects (S1-S9) and their 10 resting state sessions. The *I, J* -th block denotes the set of pairwise distances between recordings of subjects *I* and *J*. Distances are indicated by the color bar which is the same for both panels. b) Cumulative distribution functions (CDF) of intra-subject distances (solid) and inter-subject (dashed) distances for the tangent space projected FCs (blue) and regular FCs (red). c) Classifier accuracy as a function of the classification threshold *θ* for both the TS method (blue) and non-TS method (orange). The dashed-blue line denotes the accuracy of random assignment. Black line at accuracy of one is provided for visual aid. d) Receiver operating characteristic (ROC) curve for classifying intra vs inter-subject distances using the TS, non-TS, and geodesic methods. Inset: The Precision Recall (PR) curve for the same classification. e) Average distance from the first recording of each patient for both the TS (blue) and non-TS (red) methods. Both curves have been shifted down to be centered around zero to allow for comparison (originally at 0.61 for TS, 0.25 for non-TS), but the variance of the curves around zero has not been changed. Error bars denote the standard deviation across the nine subjects.

Figure 2b shows the cumulative distribution function (CDF) for all the intra (solid) and inter-block (dashed) distances separately, for both the non-TS (red) and TS)(blue) method. For either method, a threshold distance *θ* defines a decision boundary, such that FC pairs *i, j* with distance *d_i,j_ < θ* are classified as belonging to the same subject. Intra-block distances with *d_i,j_ < θ* correctly identify *i, j* with the same subject (true positives), while inter-block distances with *d_i,j_ < θ* incorrectly identify *i, j* with the same subject (false positives). In terms of the inter- and intra-subject distance CDFs, perfect subject discriminability is possible only if there exists a value *θ* for which the inter-block CDF is zero, while the intra-subject CDF is one. Generally though this is not the case, meaning that false positives (an inter-block distance with *d_i,j_ < θ*) or false negatives (an intra-block distance with *d_i,j_ > θ*) will occur.

Figure 2c shows the classification accuracy (ACC) as a function of *θ* for both the TS and non-TS methods (Eq. 6). The ACC for the non-TS method peaks at 0.95. Due to data imbalance, it is important to contrast this value against a trivial classifier that assigns FCs to subjects randomly. Because 90% of the values of **D** belong to inter-subject class, such a trivial classifier would have an ACC = 0.9, which is denoted by the dashed blue line in Fig. 2c. Conversely, the TS method peaks at 0.9998, well above the trivial classifier accuracy. The range of *θ* for which the accuracy is near unity is also much broader for the TS method than the non-TS method, implying that the TS method is more robust with respect to the specific choice of *θ*.

We also computed the Receiver Operator Characteristic (ROC) curve shown in Fig. 2d (see Methods). For the TS method the ROC appears visually constant (blue). For the non-TS method, the true positives increase monotonically with the false positives due to the overlap of the inter-intra CDFs. We use the AUC-ROC to summarize the ROC curves (see Methods). Whenever it is not explicit from the context, we will adopt subscripts *T* and *S* to distinguish between the TS and non-TS methods, respectively. We found AUC*_T_* = 0.99998 (*perfcurve* MATLAB), while AUC*_S_* = 0.94. We repeated our analysis for different regularizations, using *⋋* = 0.1 and *⋋* = 10, and observed that higher regularization improves subject discriminability. For *⋋* = 0.1, we found AUC*_T_* = 0.9997, where as for *⋋* = 10 we observe perfect subject discriminability (AUC*_T_* = 1). We also calculated the Precision-Recall (PR) curve (inset of Fig. 2d) as a complement to the accuracy and ROC (see Methods). We found the PR curve gives qualitatively similar results as the ROC curve. For the TS method we find the AUC-PR to be 0.9997, and for the non-TS method the AUC-PR is 0.98. For clarity, we will focus on the ROC curve for the remainder of the manuscript.

For comparison we calculated distances using the geodesic distance as well (Eq.2). As in (Abbas et al., 2021), we found that classification efficacy depends heavily on the regularization, and only outperforms the non-TS method for a relatively narrow range of values (see Supporting Information). Within this range we found an optimal value at *⋋* = 1.46, which is consistent with those observed by Abbas et al. (2021). For this value, the classification accuracy reaches a high of 0.96 (*θ* = 9.98), though this was not included in Fig. 2c as the range of *θ* were very different (*θ* ∈ [5, 20]). The ROC and PR curves are shown in Fig. 2d. We find an AUC-ROC of 0.97, and AUC-PR of 0.99. This suggests that for an optimized value of the regularization, the geodesic distance can perform better than the non-TS method, but we never observe it to outperform the TS method. Given this, we exclusively focus on the TS method in the following.

We also tested if there was observable drift in functional connectivity over the 10 sessions, spanning ten subsequent days. For each subject, the FC corresponding to the first session was chosen as the reference. For each of the nine subsequent sessions, the distance from the initial session was calculated. Fig. 2e) shows the mean distance averaged across subjects for both the TS and non-TS methods. Neither the TS nor non-TS methods displayed any significant trend. To do a more fine-grained analysis of potential day-to-day differences, we also subdivided recordings into non-overlapping blocks of equal size. FCs generated from blocks belonging to the same session were minimally, but significantly closer to each other than FCs belonging to different sessions (see Supporting Information Fig. S2). This is consistent with day-to-day variations in a subject’s head placement. Interestingly, only the TS method was able to observe these differences, suggesting the method has higher sensitivity.

Lastly, the above analysis compares FCs pairwise. This approach may not be representative of how fingerprinting would be used in practical setting where an unknown FC is compared against a database of labeled FCs (Abbas et al., 2021; Finn et al., 2015). Using the approach outlined in Methods, we tested the efficacy of identifying the corresponding subject of a randomly chosen unknown FC to a database of known FCs. Because of the large number of sessions in the MSC dataset, we are also able to test the effect of the size of the database. This was done by choosing *k* random FCs per subject to form the database (without replacement). The unknown FC is randomly chosen from the remaining FCs. For each *k* ≥ 1 the TS method identified the subject of the unknown FC perfectly. The non-TS method showed a dependence on the database size. For *k* = 1 the identification rate was 97%, which is on par with the classifier accuracy. As the database size increased, subject identification rates improved up until *k* = 4 where it saturated at 100%.

For each of the three tasks (motor, memory, mixed) we generate the distance matrix **D** separately. For the non-TS method these are shown in Fig. 3a. As mentioned in the Methods, several recording days and one subject had to be removed from the task-based analysis due to missing auxiliary fixation masks. We also note that the length of the recordings was not controlled for here, but is analyzed in a later section.

**Figure 3.**
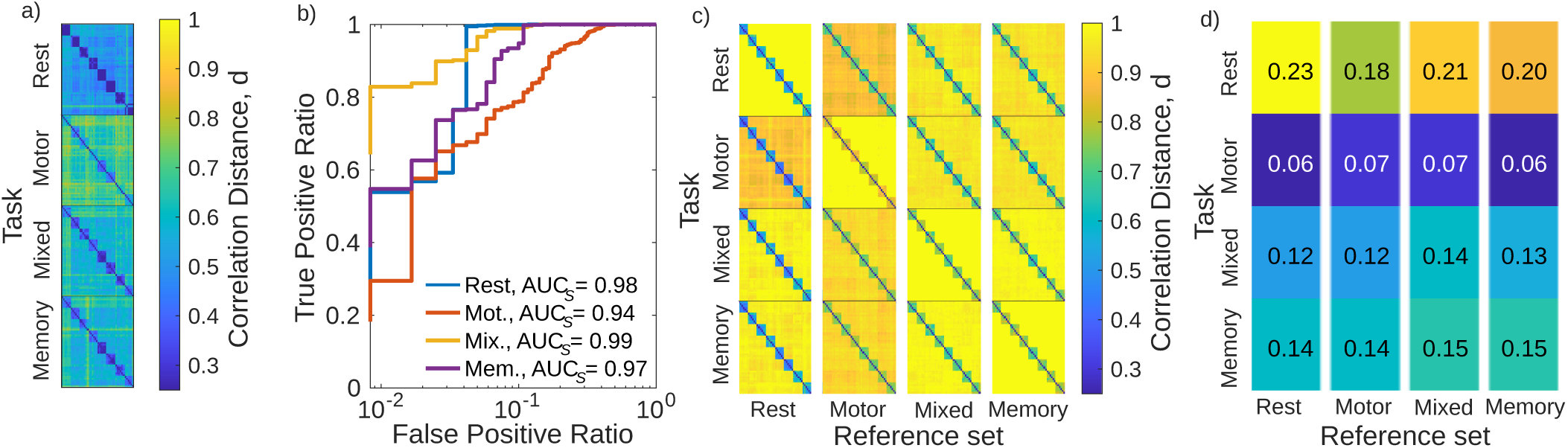
a) Distance matricies for the three tasks and rest using the non-TS method. Within each block are the correlation distances between each pair of FCs. b) The ROC curve for each of the cases shown in panel a). The corresponding AUC*_S_* is also provided. c) The distance matricies for TS method. Rows represent which category the recordings to be fingerprinted belong to. Columns represent which category the recordings that make up the tangent space reference **C**_ref_. belong to. Within each block are the correlation distances between the tangent space projected FCs, using the indicated reference. For each block AUC = 1. d) The degree of separability as measured by Δ (Eq. 9) corresponding to each case as ordered in panel c.

Figure 3b shows the ROC curves for the individual tasks when using the non-TS method. As before we quantify the ROCs using the AUC. we find that the Mixed and Memory tasks performed the best (AUC*_S_* = 0.99 and 0.97, respectively), while the motor task performs the worst (AUC*_S_* = 0.94). Due to different number of sessions and subjects, we also reanalyzed the rest case with the same sessions removed for consistency. We found that AUC*_S_* = 0.98, an increase from the 0.94 in the previous section. We suspect that this is due to the task of fingerprinting being easier when the number of subjects is decreased (Ramduny & Kelly, 2024; Waller et al., 2017).

The choice of reference point for the TS method introduces a new degree of freedom not present in the non-TS method. The first column of Fig. 3c shows the distance matricies for each task (along rows) when the reference is generated from FCs belonging to the rest case. We then changed the reference set to be that of all FCs belonging to a particular task, and repeated the analysis, which is shown in the remaining columns of Fig. 3c. In all cases AUC*_T_* = 1. To better gauge how well the identification is performing, we calculate the difference between the smallest between-subject distance and the largest in-subject distance:

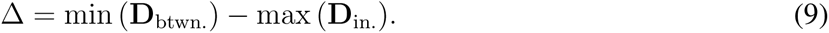

Because the AUC is one in all cases, Δ is positive for all cases, as shown in Fig. 3d. Going across columns (i.e., keeping the set of recordings to be identified the same but changing the reference), Δ appears largely constant. The most notable change is for rest recordings (top row); when using the motor-task recording as a reference there is a drop to Δ = 0.18 from the optimal Δ = 0.23 obtained when using the rest recordings as reference. Going down the rows rest recordings have the greatest Δ, motor-task the lowest, and memory/mixed-tasks are between the two. While true functional differences could play a role, we believe this more likely reflects the discrepancy in recording length (30 minutes for rest, roughly 13.5 minutes for mixed and memory tasks, and 8 minutes for motor tasks). Indeed, we test this in the Supporting Information (Fig. S3) and observe that Δ is comparable across all cases presented in Fig. 3d when accounting for the scan length.

### Discriminability of Resting-State Subnetworks

The Gordon 333 atlas is subdivided into thirteen non-overlapping resting-state subnetworks. Here we test the efficacy of subject discriminability using these subnetworks in isolation. Only subnetworks with 10 or more ROI were considered, leaving eight in total. For each subnetwork we calculate the ROC-AUC for the three task cases and all available rest recordings. ROC-AUC scores for the non-TS and TS methods are shown in panels *a* and *b* of Fig. 4, respectively. Table 1 summarizes the mean AUC averaged across all subnetworks of each case (denoted as 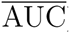).

**Figure 4.**
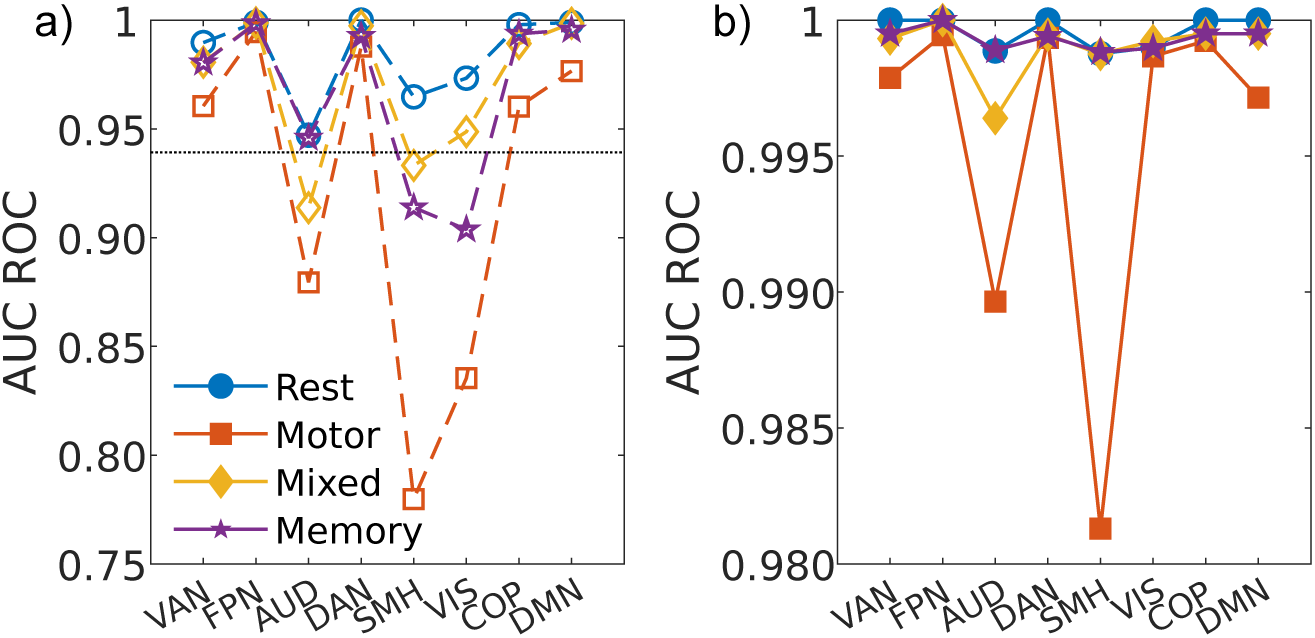
a) The area under the curve (AUC) of the receiver operating characteristic (ROC) curve for different resting-state networks in the context of rest conditions and different task conditions obtained using the non-TS method. The dotted horizontal line shows the AUC for the whole brain analysis for reference. Lines are used as visual guide. b) Same as a) but for the TS method.

**Table 1.** For TS and non-TS methods the minimum, mean (bold), and maximum (in that order) ROC-AUC scores for the different tasks, are provided. The mean value is calculated as the average of all AUC scores across all eight of the resting state subnetworks considered.

| $\overline{\text{AUC}}$ | TS | non-TS |
| --- | --- | --- |
| rest | 0.9988, **0.9996**, 1.000 | 0.8853, **0.9690**, 1.000 |
| motor | 0.9813, **0.9953**, 0.9994 | 0.6925, **0.9024**, 0.9993 |
| mixed | 0.9964, **0.9990**, 1.000 | 0.8161, **0.9306**, 0.9995 |
| memory | 0.9988, **0.9993**, 1.000 | 0.8452, **0.9406**, 0.9995 |

Focusing first on the non-TS method, we find the average ROC-AUC for the rest case to be 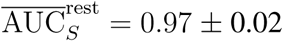 (errorbars denote one standard deviation across the different subnetworks). The AUD, SMH, and VIS subnetworks performed the worst (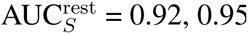, and 0.96, respectively), while FPN, DAN, and COP performed the best (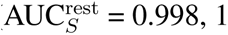, 1, and 0.995 respectively). The performance of these six subnetwork follows exactly the ‘processing’ and ’control’ subdivision proposed by (Power et al., 2011). We also find the VAN and DMN also perform well (AUC = 0.986 and 0.998, respectively), though these are not categorized as processing and control.

These trends were observable when analyzing the task based recordings as well; AUD, SMH, and VIS perform the worst (AUC*_S_* = 0.84, 0.75, and 0.80, respectively). Of the remaining subnetworks, VAN, COP, and DMN have a noticeably reduced AUC score. This is largely consistent with the results by (Finn et al., 2015), who used non-TS analysis for fingerprinting. For the mixed and memory tasks, we found 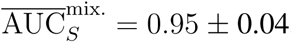, and 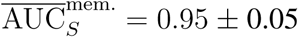. Subject discriminability was worst in the motor tasks, with 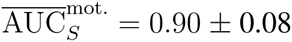.

The fingerprinting efficacy of some subnetworks was found to be task dependant while others were task independent. Generally, the ‘processing’ networks outlined above (AUD, SMH, VIS) were highly task-dependant, while the FPN and DAN were almost entirely task independent. Interestingly, while these processing nodes are often the worst performing in terms of fingerprinting efficacy, in the rest-based recordings they are still capable of out-performing the rest case whole-brain analysis (AUC*_S_* = 0.94, dashed line in Fig. 4a).

As Fig. 4b) shows, the TS method outperforms the traditional method for all subnetworks across all cases tested, never going below AUC*_T_* = 0.98. The resting and memory cases both performed strongly across all subnetworks with 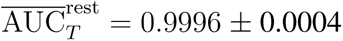 and 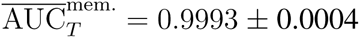. For mixed tasks, we found 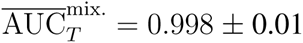, with the AUD subnetwork standing out as performing worse than all other subnetworks (AUC*_T_* = 0.995 ± 0.001). Discriminability was again lowest during motor tasks (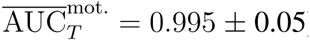). Most strikingly, while the AUD and SMH again performed the worst (as in the non-TS method), the VIS subnetwork performed comparable to the processing subnetworks, despite its categorization as a control subnetwork. It is conceivable that this reflects the TS method’s increased sensitivity. We also note that these results also did not qualitatively change for regularizations of *⋋* = 0.1 and *⋋* = 10.0, and for all values of *⋋* out-performed the non-TS method (see Fig. S4). However, in contrast to the regularization tests performed above, here we found smaller regularization fared better at subject discrimination.

Finally, we repeated the fingerprinting using the unknown target FC with a known database for the SMH subnetwork in the motor task. We chose this as it presents a worst-case scenario in terms of how large the database would need to be for fingerprinting. As in the resting case, we randomly choose *k* FCs per subject to form the database, and a random target FC is chosen from the remainder. For the TS method we observe that for *k* = 1 the identification success rate was 95%, which increases to 99% at *k* = 4 (we did not test above this due to the reduced number of sessions in the task data as explained in the methods). For the non-TS method, the success rate at *k* = 1 was 78%, which increased to 87% at *k* = 4. This suggests that for this worst-case scenario, fingerprinting requires *k >* 4, at least in the non-TS method.

### Effect of Data Length

Here we test the efficacy of FC classification as a function of data length. We temporally subsample the whole-brain recordings from which FCs are generated. A boot-strapping method is used to calculate an average AUC (see Methods for further details). Because periods of motion are removed, results are reported as the number of frames being used, though we provide what that time would have been when sampled at 2.2 Hz for reference.

First we analyzed resting state recordings which were subsampled down to between 16 and 151 frames (30 seconds to 5 minutes). Identification using the TS method was effectively independent of the recording length, saturating close to unity even for 16 frames (Fig. 5a). Conversely, classification using the non-TS method performs worse at identifying subjects for the same number of frames, with AUC*_S_* = 0.83 ± 0.02 at 16 frames. While we observe a rapid increase to AUC*_S_* = 0.968 ± 0.001 until 46 frames (roughly 1.5 minutes), the AUC*_S_* does not saturate to 0.98 until approximately 100 frames (3.5 minutes).

**Figure 5.**
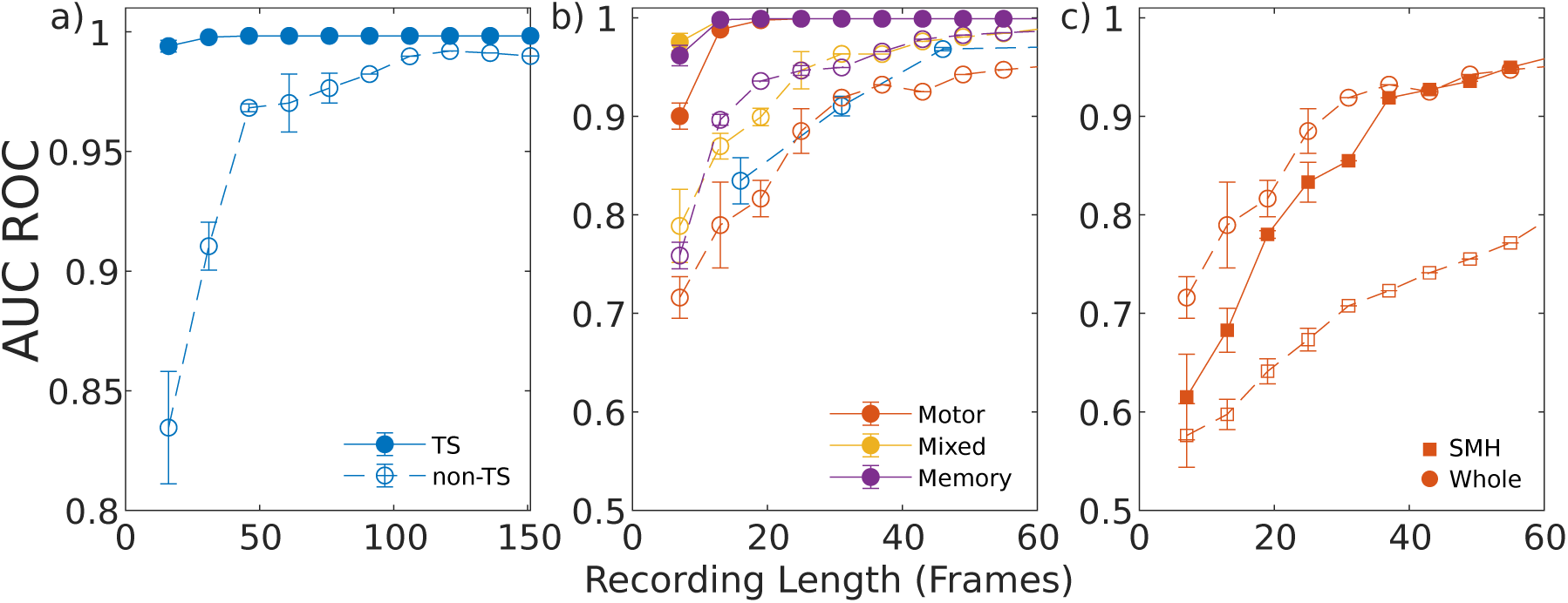
The mean AUC-ROC values for different recording durations, averaged over all bootstrap samples. Across all three panels, solid markers denote the TS method, with hollow markers for non-TS. Lines shown for visual aid. The following cases are considered: a) Resting state recordings across 9 subjects and 10 sessions. b) Task based recordings. The resting state non-TS method (blue) is included for comparison. c) The SMH resting state network during motor task (square markers) for the TS and non-TS methods. The whole-brain motor task using the non-TS method is included for reference (circle). Error bars denote one standard deviation across all bootstrapped samples. To avoid oversampling, the number of samples is inversely related to the recording length, which results in smaller error bars for the larger recording lengths.

Task recordings were subsampled to between 7 and 61 frames (15 seconds to 2.2 minutes) due to their shorter original duration. Fig. 5b) shows that the TS method is consistently near unity, though for the shortest recording lengths we do observe a decrease in the AUC*_T_*, especially for the motor task (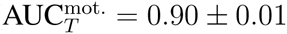). Again the non-TS method performs poorly with the smallest segments (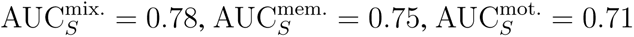), followed by a rapid rise until 40 frames, and a slow increase there-after, which is consistent with Fig. 5a. We also observe the motor tasks perform the worst across all recording durations, with 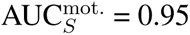 for the longest segments. Fig. 5b) also shows that, when controlled for length, the mixed and memory tasks outperform the resting case using the non-TS method, whereas the motor task does not.

Finally, we tested the effect of data length on the resting-state subnetworks of the Gordon 333 parcellation. We focused on the SMH subnetwork in the motor task, as this was the worst performing subnetwork across all cases. The results are shown in Fig. 5c. For the shortest segments, both TS and non-TS methods perform poorly at subject discrimination, with 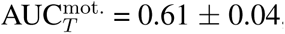, and 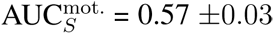 (respectively), both of which are worse than the whole brain analysis and comparable to chance levels (AUC = 0.5). While discriminability using the AUC*_T_* improves to 0.96 for 60 frames, AUC*_S_* only reaches 0.78 for the same amount of data, though it does not appear to yet be saturating.

## DISCUSSION

We performed functional connectome fingerprinting of resting-state and task-based fMRI recordings belonging the midnight scan club dataset. We applied two approaches to performing this task – a geometry-aware tangent space method which takes into account the curvature of the positive-semidefinite matrix space to which functional connectomes belong, as well as the standard method of calculating correlation distances directly form FCs, which does not take this curvature into account. In each analysis we compared the efficacy of subject identification of both methods.

We first performed classification of the 90 resting state recordings (10 recordings per subject) of the MSC. Consistent with the fingerprinting analysis performed in (Finn et al., 2015) we observed that, for a given discrimination threshold *θ*, the non-TS method was capable of reaching a high accuracy of subject discrimination (ACC = 0.95). However, the range of *θ* for which the method outperformed a trivial classifier (e.g., one that randomly assigns FCs) was small. Conversely, the TS method outperformed the non-TS method in all measures of classifier efficacy (accuracy, ROC, and PR), achieving near perfect scores in each, and over a considerably wider range of *θ*. This latter aspect is particularly important for expanding fingerprinting analysis to practical settings – a wide range of possible discrimination thresholds would allow for the addition of new data without the need to recalculate an optimal identification threshold.

One important question with regards to subject identification using FCs is whether the method simply reflects anatomical differences which could artificially improve fingerprinting efficacy (Gordon, Laumann, Adeyemo, & Petersen, 2017). Our analysis showed that subject discrimination efficacy is state dependent. This was especially true for the analysis of resting state subnetworks where certain subnetworks displayed strong task dependence while others did not. Thus, while anatomical differences may contribute towards subject discriminability, we believe these observations cannot be explained by this alone. Likewise, previous studies on FC fingerprinting (using the non-TS method) address this by applying a smoothing kernel to the BOLD signals prior to registration to a common atlas, finding this led to only a small decrease in fingerprinting efficacy (Finn et al., 2015), further suggesting evidence that the method reflects more than anatomical differences.

The above point is important as the clinical usefulness of fingerprinting methods may lie in longitudinal observations of a single subject, instead of discriminating between multiple subjects, where anatomical changes would be expected to be minimal (or at least on shorter time scales). For example, FCs have been applied to study network changes in the context of neurological diseases and disorders such as major depressive disorder (MDD) (Supekar et al., 2008; Wang et al., 2012). While MDD is associated with anatomical differences relative to healthy controls (Korgaonkar, Fornito, Williams, & Grieve, 2014), FCs could be used to track the efficacy of treatment on shorter time scales, where significant anatomical changes would not be expected. Indeed, recently the fingerprinting approach was used to study treatment response and predictors of treatment response for MDD (Nemati et al., 2020), however without the TS method. It is possible the increased sensitivity of the TS method would potentially prove a superior biomarker (relative to the standard non-TS method) in FC studies of not only MDD, but also potentially more general brain disorders (Du et al., 2018; Kaufmann et al., 2018).

Expanding on clinical applicability, we also studied how the size of a pre-existing database can effect fingerprinting efficacy when a new FC is introduced. Prior studies have focused on datasets containing test-retest data which only contained two FCs, limiting the extent to which this question can be studied (Finn et al., 2015). For the non-TS method we found that fingerprinting efficacy improves monotonically with the database size, eventually saturating at perfect discriminability. For the resting state data this was found to occur at four FCs, however the exact point depends on the task, and may also depend on the duration of the recordings. Conversely, the TS method performs fingerprinting perfectly with only a single FC per subject in the database for resting-state data. Furthermore, we observed that the TS method was capable of near-perfect classification after only a minute worth of scans, while the non-TS method could take three minutes worth and would still not match the TS method in terms of identification efficacy. This resource efficiency could be an important point in pediatric research where frequent in-scanner movement poses a great challenge to data aquisition as well as the comfort of the patient (Ellis et al., 2020). The significantly lower amount of data required by the TS method could shorten the acquisition time required, and when paired with other methods (e.g., movie-viewing (Vanderwal et al., 2017)) could ease the overall procedure, while still maintaining the usefulness of existing data. We note, however, that an important limitation in this study is the number of subjects (nominally 9), which certainly leads to higher classification efficacy (Ramduny & Kelly, 2024; Waller et al., 2017). While we conjecture that the TS method would still outperform the non-TS method with more subjects, the rate of potential efficacy decrease as a function of the number of subjects, especially relative to the non-TS method, is not known and is not addressable using the MSC data alone. Tackling this remains an interesting challenge for the future.

We also tested subject discriminability in the context of task-based recordings, which included a motor task, incidental memory, and a mixed design task. Using the non-TS method we observed that, when differences in data length were controlled for, subject discrimination improved for mixed and memory tasks relative to rest. This is largely consistent with what was also observed by Abbas et al. (2023), except that we observed motor tasks perform worse relative to rest. Whether this is related to the differences in the tasks (Rai, Graff, Tansey, & Bray, 2024) performed (detailed in (Barch et al., 2013)), would be interesting to investigate in the future. Despite this, it is interesting to observe this state-dependant subject discriminability and how it may relate to the task. The work presented in (Abbas et al., 2023) proposes a “fingerprinting gradient” in the context of decreasing subject discrimination efficacy from individuals to mono-zygotic twins and finally dizygotic twins. Perhaps it is possible to postulate a conceptually similar fingerprinting gradient going from “cognitively heavy” tasks (mixed and memory), down to motor tasks which involve more stereotyped neural activity, with resting state potentially being a boundary between the two. This remains an exciting challenge for the future. However, when using the TS method, all tasks allowed for near-perfect subject discrimination, so long as the scan was at least around a minute in length. Interestingly, we observed no significant dependence on the choice in reference (e.g., using a reference generated from motor-task FCs to fingerprint resting-state FCs). This is useful as it means that a database of rest recordings (as an example) can still perform well in discriminating task-derived FCs.

While subject discrimination performed well on the whole brain level, fingerprinting using resting-state subnetworks also performed as well, and sometimes better. We find the efficacy of a subnetworks ability to perform fingerprinting aligns closely with its membership to the “control” or sensory/motor “processing” networks (Power et al., 2011). The control network, containing the FPN, DAN, and COP, is thought to be more flexible in order to be able to adapt to novel tasks (Zanto & Gazzaley, 2013). Other studies have also found that the FPN, VAN, DMN, and DAN exhibit high inter-subject variability, all of which were found to have the greatest performance in our analysis (Mueller et al., 2013; Vanderwal et al., 2017). We speculate this flexibility and variability is responsible for subject discrimination, an observation which has also been reported in (Finn et al., 2015). These subnetworks, especially the FPN, are also thought to have hub-like properties in relation to various tasks, which could explain why fingerprinting efficacy was unchanged by tasks (true for both TS and non-TS methods) (Cole et al., 2013). Likewise, while not necessarily a member of the control network, the DMN interacts strongly with the FPN (Elton & Gao, 2014) and is involved in cognition, motivating its high fingerprinting efficacy.

Conversely, the more static nature of processing networks (VIS, AUD, SMH) (Zanto & Gazzaley, 2013) could be related to their poor performance at fingerprinting, at least when using non-TS methods. Other studies have also found that inter-subject variability was lowest in these unimodal sensory subnetworks (Mueller et al., 2013; Vanderwal et al., 2017). While these subnetworks performed poorly relative to other subnetworks when using the TS method, objectively speaking their AUC scores were high. One striking exception was the visual subnetwork (VIS); using the non-TS method the VIS subnetwork performed in line with the other processing networks (i.e., poorly), but using the TS method it performed comparable to processing networks and was task independent. Of course, visual stimuli and feedback are important aspects of motor tasks, memory, and cognition in general (Cavanagh, 2011; Hofstetter, Achaibou, & Vuilleumier, 2012) which could motivate this finding, but does not on its own explain why the AUD network does not display the same. Visual memory has been demonstrated to be stronger than auditory memory (Cohen, Horowitz, & Wolfe, 2009), which could be related, but this is speculative. If this observation is reproducible in other studies, we believe this would form an interesting point of investigation from both the neuroscience and mathematical point of view.

While control networks were state-independent, the fingerprinting efficacy of processing subnetworks was not (with the aforementioned exception). Efficacy was lowest during motor tasks, ubiquitously improving in the rest state and even in the mixed and memory tasks, and often outperformed the whole-brain FCs (at least, for the non-TS method). This brings up two points: First, the reason why process subnetworks can so drastically improve their fingerprinting efficacy during rest is not entirely clear. It could be related, for example, to the FPN and executive control (see above), or distributed coding of neural activity (Steinmetz, Zatka-Haas, Carandini, & Harris, 2019). Second, the fact that subject discriminability improves using subnetworks could suggest that the various subnetworks cause a blurring effect in whole-brain FCs – perhaps from the processing networks themselves. This effect could be useful to consider when studying the early stages of disease when differences may be less obvious, such as in the use of FCs to study Alzheimer’s disease (Supekar et al., 2008). Moreover, we believe the variation in fingerprinting efficacy with region and task (and the dichotomy between the ‘control’ and ‘processing’ areas) further suggest that inter-subject functional differences are being measured, and not just anatomical ones.

The distances between FCs can provide information about the intrinsic differences of individual functional brain dynamics. While comparing FCs between individuals can offer interesting insights, its potential use for understanding the evolution of a single individuals brain dynamics longitudinally, such as management of disorders or disease, is perhaps even more exciting. The results above motivate that geometry aware methods (like tangent space analysis) can utilize both new and existing data more efficiently, and with greater accuracy.

## ACKNOWLEDGMENTS

This project was supported by Alberta Innovates (232403074). J.D. was supported by the Natural Sciences and Engineering Research Council of Canada (RGPIN/05221-2020). S.K.V.K. was supported by Mitacs through their Globalink Research Internship program.

## TECHNICAL TERMS

**Functional Connectome/ Connectivity (FC)** A matrix representation of the brain connectivity derived from the pairwise correlations of brain dynamics (such as those recorded by fMRI).

**Fingerprinting** The procedure of identifying the subject(s) that produced an unknown FC(s).

**Positive Semi-Definite** A mathematical property that all eigenvalues of the FC are greater (or equal to) than zero.

**Positive Semi-Definite Manifold** The mathematical space of positive semi-definite matricies, which is a sub-space of all possible matricies,

**Compact Convex Manifold** A space that is does not extend infinitely and has no holes, and whose curvature is convex everywhere. An example of such a space would be a sphere.

**Geometry-aware Methods** Approaches to calculating distances between two FCs that account for the curvature of the space.

**Tangent Space** A particular geometry-aware method which approximates distances by finding a flat space that is tangent to a point of a curved space. Analogous to observing the Earth to be flat locally, despite its convex curvature.

**Reference Matrix** The point at which the tangent space is calculated, calculated here as the logarithmic average of all FCs.

**Regularization** A mathematical procedure of adding a positive value *⋋* to the diagonal of the FC to ensure it is inevitable.

**Receiver Operator Characteristic (ROC)** A plot of the true positive rate and false positive rate used to visualize the performance of a classifier.

**Precision-Recall (PR)** A plot of the relevant items among the returned items (precision), against the relevant items that were actually returned (recall).

**Area Under the Curve (AUC)** A commonly used metric for summarizing the ROC or PR curves.

