## supplemental material for "Efficacy of functional connectome fingerprinting using tangent-space brain networks"

*Journal:* **NETWORK NEUROSCIENCE**

RESEARCH

#### *Classifier efficacy of geodesic distance depends on regularization*

The geodesic distance between two positive semi-definite matricies $\mathbf{P}_{1,2}$ is given by (Bhatia, 2015):

$$\delta(\mathbf{P}_1, \mathbf{P}_2) = \left(\mathrm{Trace}(\log^2_{\mathrm{m}}(\mathbf{P}_1^{-\frac{1}{2}}\ \mathbf{P}_2\ \mathbf{P}_1^{-\frac{1}{2}}))\right)^{1/2}. \quad (1)$$

Unlike the TS method, the geodesic distance satisfies the requirements of a true distance metric (including symmetry). However, the geodesic distance requires $\mathbf{P}_1$ to be invertible, which may not always be the case. As discussed in the manuscript, one remedy is to include a regularization term $\mathbf{P}_1 + \lambda\mathbf{1}$ (Venkatesh, Jaja, & Pessoa, 2020). In classification tasks, it has been shown that efficacy may improve with regularization even when it is not technically needed, and there is an optimal range of $\lambda$ (Abbas et al., 2021). Fig. S.1a shows the ROC-AUC for classification of the rest case using the geodesic distance as a function of $\lambda$. We also include the TS method and non-TS (recall however the non-TS method does not use regularization). We note first that the AUC for the geodesic distance is much more dependent on regularization than the TS method which is essentially independent of it. Second, the geodesic distance outperforms the non-TS method, but only in a narrow window of $\lambda$ between roughly 0.6 to 8, and even then only marginally so.

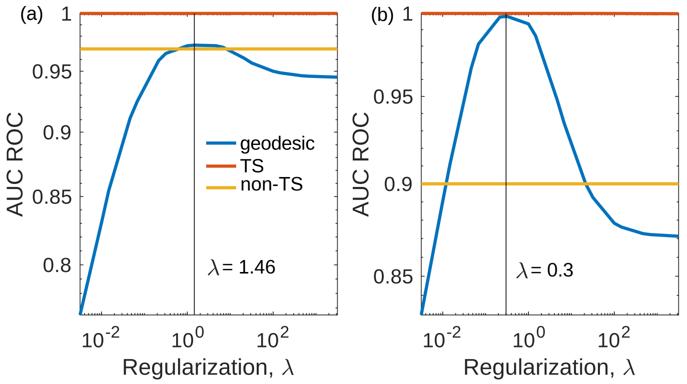


**Figure S.1.** a) The ROC-AUC of subject descriminability in the resting state for the geodesic distance, tangent space (TS), and non-TS method, as a function of the regularization parameter $\lambda$. The vertical line denotes the optimal value of $\lambda$ for the geodesic distance. Note that while the non-TS method is truly independent of regularization, the TS method does indeed vary, but is not discernable at this scale. b) Same as previous panel but for the motor-task set of recordings.

We also tested the motor task recordings as these were the worst performing. Interestingly, here we see the geodesic distance outperform the non-TS method for a much wider range of $\lambda$, and also attains much higher values as well. Why this would be the case is currently not clear however. Regardless, in no case do we observe the geodesic distance outperform the tangent space method. This, paired with the high dependence on the regularization (not to mention the actual decision boundary $\theta$) suggests the tangent space method as superior to the geodesic distance in tasks of subject classification.

#### *Comparing subdivisions of recordings*

To test for day-to-day differences we subdivide recordings into smaller segments, recalculate FCs for each subdivision, and then find the distances between FCs. In greater detail; we subdivided each rest session into blocks of 150 frames (approximately five minutes). Due to motion artifact removal the exact number of resulting blocks varied between three and four per recording. For each block we calculated the FC, meaning each session had three to four FCs per session. Initially we compared the distances of the FC belonging to the first block against all remaining FCs. For clarity we will denote this FC as $FC_0$. For the TS method $FC_0$ also serves as $\mathbf{C}_{\mathrm{ref.}}$. While FCs belonging to the same group as $FC_0$ (the in-group) were indeed closer then FCs of later blocks (the out-group), this approach suffers from skewed statistics as there are nine times as many out-group blocks as there are in-group blocks. As we did not observe any differences in the distance metric beyond the second session, we split the recordings according to odd and even numbered recording days and compared in pairs. Specifically, $FC_0$ corresponds to the first block of every odd-numbered session. In- and out-group distances were then compared as before, this time restricting the out-group to be only the FCs from the subsequent (even-numbered) session. This ensured balanced statistics between both the in and out group (122 and 124, respectively). Finally, a two-sample Kolmogorov-Smirnov test (*kstest2*, MATLAB) was conducted to test if the resulting distributions of in-group and out-group distances were significantly different.

Fig. S.2 shows box plots for both the TS and non-TS methods. For the TS method the in group is found to have smaller distances from $FC_0$ relative to the out-group, as expected. The median distance for the in-group was 0.56 (0.55,0.58 for upper and lower quartiles) and 0.61 (0.60,0.63) for the out-group. The KS test rejects the hypothesis that the in- and out-groups were obtained from the same distribution ($p = 10^{-20}$). Conversely, using the non-TS method, the in- and out-groups show no significant difference, with a median of 0.66 (0.63,0.69) for the in-group and 0.65 (0.62, 0.69) for the out-group. The KS test does not reject the hypothesis that these two were obtained from the same distribution ($p = 0.589$). This result suggests that the TS method is sensitive enough to discern day-to-day differences between FCs. However, our analysis does not allow us to definitively determine if these day-to-day differences are due to true changes in FC or if they are due to variability in scanning (e.g., head position).

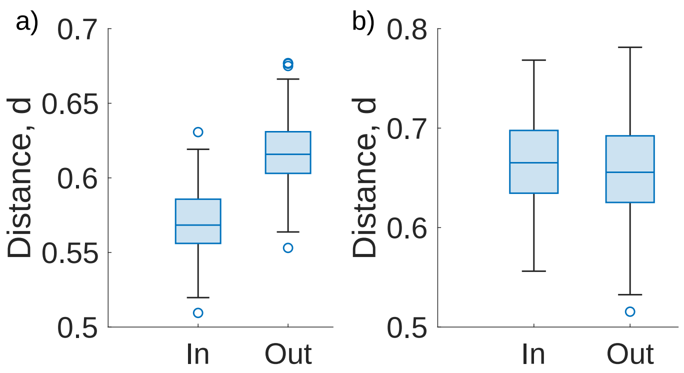


**Figure S.2.** Here we subdivide each recording into blocks of equal length and generate FCs for each. Choosing one FC as reference, box plot shows the distance calculated using the TS method (panel a) and non-TS method (panel b) between FCs blocks belonging to the same recording (in-group) and blocks belonging to the subsequent recording session (out-group). Box edges represent quartiles (with median line shown), with whiskers denoting maximum and minimum.

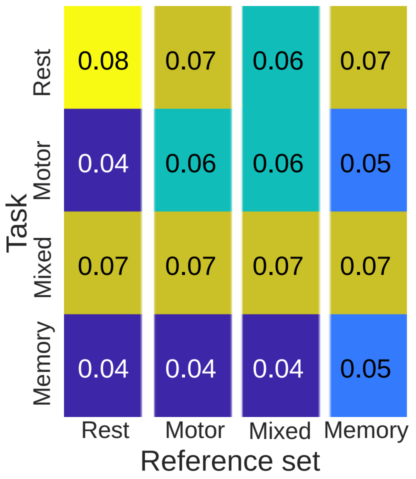


**Figure S.3.** Same as Fig. 3d but when each recording is of equal length (3 minutes). Each block indicates the $\Delta$ (Eq. **??**) seperation between the intra- and inter-subject CDFs. Rows represent which category the recordings to be fingerprinted belong to. Columns represent which category the recordings that make up the tangent space reference $\mathbf{C}_{\mathrm{ref.}}$ belong to. Within each block are the correlation distances between the tangent space projected FCs, using the indicated reference.

#### *Reference Set With Controlled Length*

In Fig. 3d we study the discriminability of FCs belonging to the different task categories, as well as the effect of changing the reference point. Here we repeat the analysis but control for the length of the recording. This was done by taking the first 85 frames of each scan. This value was chosen as it was the duration of the shortest scan post-motion artifact and non-fixation removal. We repeat the TS method as in the main text and observe AUC = 1 in each case (consistent with the recording duration analysis in the main text). We calculate the intra- and inter-subject distance CDF seperation using $\Delta$ (Eq. 9). Fig. S.3 shows that once scan length is controled for, the seperability of FCs is comparable across all cases.

#### *Regularization analysis for subnetworks*

In Fig. S.4 we show the subnetwork analysis repeated for $\lambda = 0.1$ and $\lambda = 10$. Interestingly, here we find that smaller regularization appears to improve discrimination rates, with over-all AUC scores improving as $\lambda$ decreases.

### REFERENCES

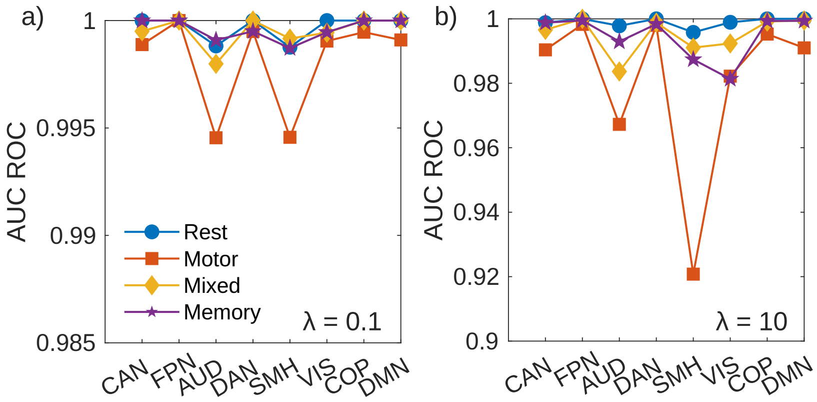


**Figure S.4.** AUC-ROC scores for the different resting-state subnetworks using regularization of $\lambda = 0.1$ (a) and $\lambda = 10$ (b).

Abbas, K., Liu, M., Venkatesh, M., Amico, E., Kaplan, A. D., Ventresca, M., . . . Goñi, J. (2021). Geodesic distance on optimally regularized functional connectomes uncovers individual fingerprints. *Brain connectivity*, *11*(5), 333–348. doi: https://doi.org/10.1089/brain.2020.0881

Bhatia, R. (2015). *Positive definite matrices*. USA: Princeton University Press.

Venkatesh, M., Jaja, J., & Pessoa, L. (2020). Comparing functional connectivity matrices: A geometry-aware approach applied to participant identification. *NeuroImage*, *207*, 116398. doi: https://doi.org/10.1016/j.neuroimage.2019.116398
